# Reduced phenotypic plasticity evolves in less predictable environments

**DOI:** 10.1101/2020.07.07.191320

**Authors:** Christelle Leung, Marie Rescan, Daphné Grulois, Luis-Miguel Chevin

**Author notes:** Christelle Leung, Marie Rescan, Daphné Grulois, Luis-Miguel Chevin. **Corresponding Author:** Christelle Leung or Luis-Miguel Chevin, Centre d’Écologie Fonctionnelle et Évolutive, 1919 route de Mende, 34293 Montpellier Cedex 5, France, or.

## Abstract

Phenotypic plasticity is a prominent mechanism for coping with variable environments, and a key determinant of extinction risk. Evolutionary theory predicts that phenotypic plasticity should evolve to lower levels in environments that fluctuate less predictably, because they induce mismatches between plastic responses and selective pressures. However this prediction is difficult to test in nature, where environmental predictability is not controlled. Here, we exposed 32 lines of the halotolerant microalga *Dunaliella salina* to ecologically realistic, randomly fluctuating salinity, with varying levels of predictability, for 500 generations. We found that morphological plasticity evolved to lower levels in lines that experienced less predictable environments. Evolution of plasticity mostly concerned phases with slow population growth, rather than the exponential phase where microbes are typically phenotyped. This study underlines that long-term experiments with complex patterns of environmental change are needed to test theories about population responses to altered environmental predictability, as currently observed under climate change.

## Introduction

Phenotypic plasticity, the ability of a given genotype to produce alternative phenotypes depending on its environment of development or expression, is a major mechanism for responding to environmental variation across the tree of life (Scheiner 1993; Schlichting & Pigliucci 1998; West-Eberhard 2003). In recent years, the study of phenotypic plasticity has gained prominence in evolutionary ecology, with the realization that it contributes a substantial part to observed phenotypic change in the wild, notably in response to climate change (Gienapp *et al*. 2008; Merilä & Hendry 2014). Because phenotypic change is generally a strong determinant of population dynamics (Pelletier *et al*. 2007; Ozgul *et al*. 2010; Ellner *et al*. 2011), this implies that phenotypic plasticity can strongly impact population growth and extinction risk in a rapidly changing world, to an extent that depends on the rate and pattern of environmental change (Chevin *et al*. 2010; Reed *et al*. 2010; Chevin *et al*. 2013b; Vedder *et al*. 2013; Ashander *et al*. 2016; Phillimore *et al*. 2016).

However, despite the ubiquity and ecological importance of phenotypic plasticity, proving its adaptiveness for any particular trait and organism is particularly challenging (Ghalambor *et al*. 2007). Most evidence that plasticity is adaptive is instead indirect, for instance through comparison of the direction of plastic *vs* evolved responses to a novel environment (Ghalambor *et al*. 2015), except for rare studies where plastic responses have been genetically engineered to compare the fitness of plastic *vs* non-plastic genotypes across environments (Dudley & Schmitt 1996). However, beyond the putative advantage of being plastic, a more meaningful and quantitative question is whether a given degree of plasticity is adaptive. This question was thoroughly addressed theoretically, and the predictability of environmental variation was identified as a key determinant of the adaptiveness of plasticity, and driver of its long-term evolution (Gavrilets & Scheiner 1993; de Jong 1999; Lande 2009; Reed *et al*. 2010; Botero *et al*. 2015; Tufto 2015). In particular, theory predicts that phenotypic plasticity should evolve to lower levels in environments that fluctuate less predictably, because this leads to plastic responses that do not match future selective pressures (Gavrilets & Scheiner 1993; de Jong 1999; Lande 2009; Botero *et al*. 2015; Tufto 2015). Nevertheless, these predictions still largely lack direct empirical evidence (but see Scheiner & Yampolsky 1998; Dey *et al*. 2016), owing to the difficulty in manipulating the variability and predictability of the environment over evolutionary times. In addition, multiple independent replicates of environmental fluctuations are needed in order to account for their inherent randomness (environmental stochasticity), but this is difficult to achieve in nature.

A useful alternative to circumvent these limitations is to perform long-term laboratory experiments under controlled, yet ecologically realistic, patterns of environmental fluctuations. This approach, which was previously advocated by Chevin *et al*. (2013a), allows controlling for the level of environmental predictability, with all other things being equal - including the mean and variance of the environment -, and also permits sufficiently large duration and replication to unambiguously observe evolutionary responses to stochastic environments. Here we have applied this approach with the unicellular microalga *Dunaliella salina*, the main primary producer in hypersaline environments such as continental salt lakes, coastal lagoons, and salterns (Oren 2005; Ben-Amotz *et al*. 2009). The cell shape and content of this microalgae are known to respond plastically to salinity over different timescales, allowing for both immediate morphological responses to sudden osmotic stress, and slower physiological adjustments involving the production of metabolites (glycerol, carotene) inside the cell (Oren 2005; Ben-Amotz *et al*. 2009). Several of these traits can be measured at the individual level and at high-throughput, pushing work on phenotypic plasticity towards the realm of phenomics (Houle *et al*. 2010; Pendergrass *et al*. 2013; Yvert *et al*. 2013). Experimental evolution has been successfully performed previously with the closely related species *Dunaliella tertiolecta* (Malerba *et al*. 2018). Furthermore, we have recently shown that phenotypic plasticity plays a crucial role in the population dynamics and extinction risk of *Dunaliella salina* in a randomly fluctuating environment (Rescan *et al*. 2020). This makes *D. salina* particularly well-suited to investigate experimentally the evolution of phenotypic plasticity in response to environmental predictability.

## Material and Methods

### Experimental evolution

We followed up on the experimental evolution protocol initiated by Rescan *et al*. (2020), which we pursued for several hundred generations. Briefly, we used two genetically related strains (CCAP 19/12 and CCAP 19/15) of the halophilic unicellular microalga *Dunaliella salina*, which can tolerate a broad range of salinities. Long-term experimental evolution occurred from August 2017 to January 2019, during which we exposed 32 populations to randomly fluctuating salinity, and 4 populations to a constant intermediate salinity ([NaCl] = 2.4 M). Salinity changes occurred twice a week, by 20% dilution into 800 μl of fresh medium to account for population extinction (Rescan *et al*. 2020), using a liquid-handling robot (Biomek NXP Span-8; Beckman Coulter). At each transfer, the target salinity was achieved by mixing the required volumes of hypo- ([NaCl] = 0M) and hyper- ([NaCl] = 4.8M) saline media, after accounting for dilution of the pre-transfer salinity (Rescan *et al*. 2020). We totalized at least 139 salinity transfers and more than 500 generations (assuming ~1 generation per day (Ben-Amotz *et al*. 2009)). The populations in fluctuating environments were subjected to independent random time series over a continuous range (first-order autoregressive process, AR1), rather than a more artificial alternation of low vs high salinity treatments. All the lines had the same long-term stationary mean (*μ* = 2.4M [NaCl]) and variance (*σ* = 1) of salinity, but they differed in how salinity at a given time depends on the previous salinity, prior the last transfer, as determined by the temporal autocorrelation *ρ* of salinity (Fig. 1A & Fig. 1B) (Rescan *et al*. 2020). The predictability of environmental change upon these transfers depended on *ρ^2^*, the proportion of the temporal variance in salinity explained by the previous salinity (Fig. 1B). There were four autocorrelation treatments, for which the expected autocorrelations (over infinite time) were 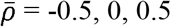, and 0.9 (Fig. 1; see Rescan *et al*. (2020) for detailed protocol), but the realized autocorrelation over the duration of the experiment spanned a continuum of values. All populations were cultivated in 1.1ml of 96-deepwell plates (Axygen^®^; Corning Life Sciences) in an artificial seawater prepared as described in Rescan *et al*. (2020), at constant temperature (24°C) and 12/12 lighting (200 μmol.m^−2^.s^−1^).

**Fig. 1.**
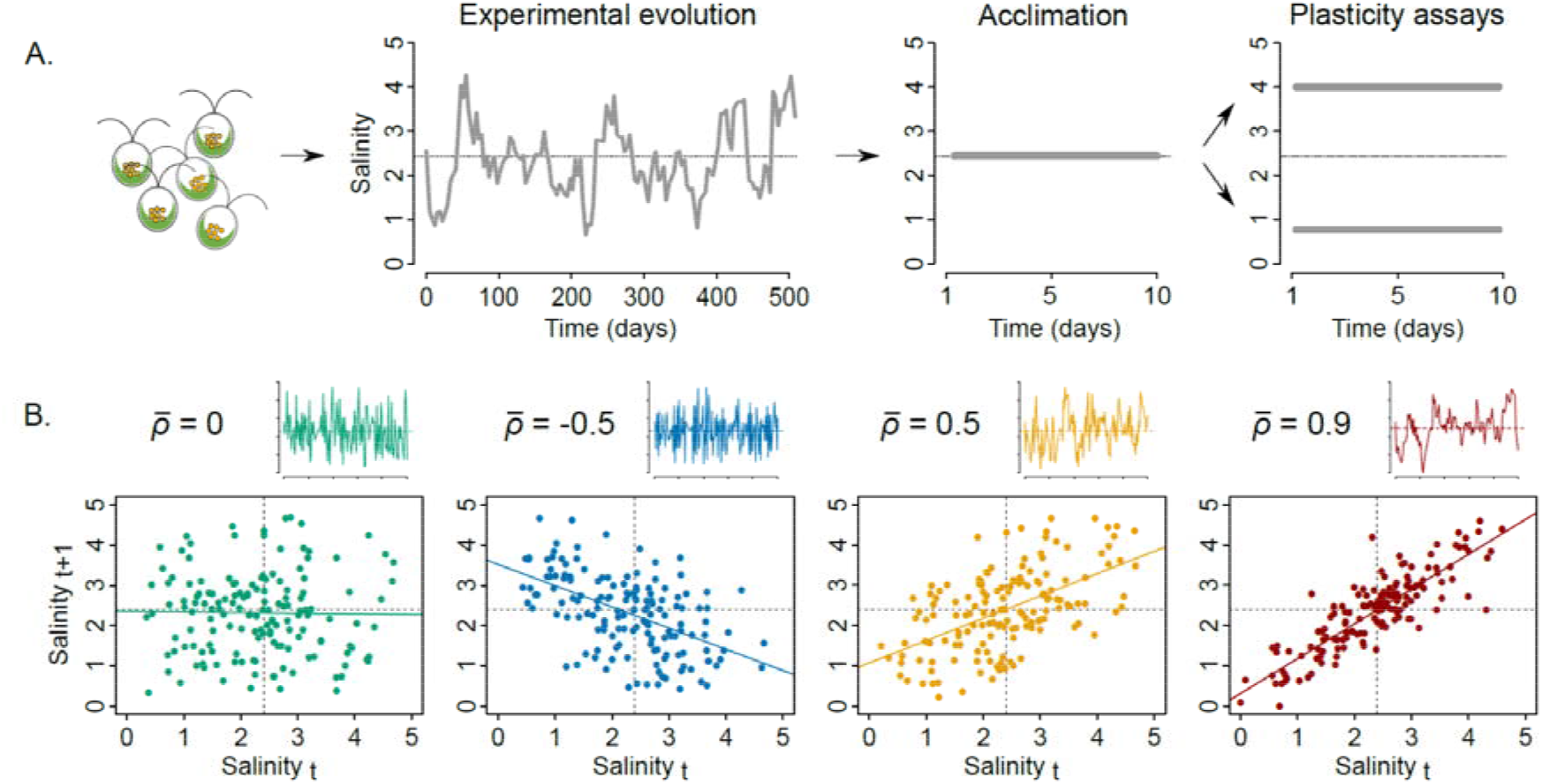
Long-term evolution experiment under variable environmental predictability. A. The different steps of the experiment are illustrated. B. Examples of the four categories of expected long-term temporal autocorrelation (). For each category, the small graph represents the associated time series for a given population (same axes as in Fig. 1A), with mean salinity shown as horizontal dotted line. The larger graphs represent the relationship between subsequent salinities in these time series. Note that the expected autocorrelations = −0.5 and = 0.5 lead to different magnitudes of salinity changes, but display the same predictability of environmental changes (^*2*^ = 0.25).

At the end of the experiment, we genetically confirmed that strains were not cross-contaminated after ~1.5 year of experiment. Specifically, for each evolved lines, we amplified one mitochondrial (333 pb fragment using the primers DsMt1-For [5’- GGTTAGTCATAGTTGGAGGT-3’] and DsMt1_-Rev [5’- GAAAACCTAACATGGCTAAGC-3’] and chloroplast (372 bp fragment using the primers DsChl1-For [5’-TTTAGGCGAATCCATAAGAG-3’] and DsChl1-Rev [5’- CCAAGCAGGTGAATTAGCTTTG-3’]) locus, specific to CCAP 19/12 and CCAP 19/15, respectively.

### Phenotypic plasticity assays

To assess the plasticity degree of the 36 evolved lineages, we compared individual cell morphologies between two salinities near the extremes of their historical range ([NaCl] = 0.8M and 4.0M). These lineages started with potentially genetically diverse strains, and were certainly also polymorphic at the end of experimental evolution. To verify that morphological changes between salinities over the 10-day assay were due to plastic responses rather than environment-specific selection, we also performed the phenotypic plasticity assays with 20 additional isogenic populations derived from five heterogenic experimental lines. One of these experimental lines was the CCAP 19/12 strain that evolved in constant salinity (2.4 M) and four others were from the CCAP 19/15 strain that evolved in constant (2.4 M) and fluctuating (with targeted autocorrelation *ρ* = −0.5, *ρ* = 0 and *ρ* = 0.9) salinities. For each of these five experimental populations, we founded four different population from single cells isolated using cells-sorting flow cytometry (BD *FACSAria*™ IIu; Biosciences-US). Because *Dunaliella salina* is haploid, we expected all cells to be genetically identical in populations founded from a single one.

The 36 lines were sampled from different salinities and at different population sizes at the end of the evolutionary experiment (Rescan *et al*. 2020). To ensure that all cells were in similar physiological states and at similar population densities at the beginning of the phenotypic plasticity assay, we first acclimated them for 10 days in the same environmental conditions. We transferred 400 μl of each experimental population into 50 ml flasks (CellStar^®^; Greiner BioOne) with 25 ml of fresh medium at salinity [NaCl] = 2.4 M, temperature 24°C, and light intensity 200 μmol m^−2^ s^−1^ for 12:12 h light/dark cycles.

Following the acclimation step, we inoculated ~2 × 10^4^ cells.ml^−1^ of each population into low (0.8 M) and high (4.0 M) salinity medium (Fig. 1), to track their population density and morphological traits for 10 days. For 24 populations (including 14 randomly chosen evolved lines and 10 isogenic populations), we also inoculated the same number of cells into 2.4 M salinity medium to assess morphological changes independent from salinity changes. We randomly placed all conditions in 2 ml 96-deepwell plates at 24°C and 12/12 lighting.

When placed in a new salinity, *D. salina* undergoes changes in cell shape and content, reflecting an osmotic adaptability that unfolds over different timescales (Ben-Amotz *et al*. 2009). Rapid changes in cell volume in response to changes in extracellular osmolarity are made possible by lack of a rigid cell wall. This then triggers longer physiological responses involving the production of osmolites (especially glycerol), changes in gene expression, and the production of diverse salt-induced proteins (Azachi *et al*. 2002; Oren 2005; Ben-Amotz *et al*. 2009), which all modify the cell content. To assess the changes in cell morphology, we passed a 150 μL sample of each population through flow cytometry at 11 time points: at the end of the acclimation step (day = 0), 4h after environmental changes (day = 1), and once per day from day = 2 to day = 10.

Intrinsic structural parameters of cells were measured using a Guava^®^ EasyCyte™ HT cytometer (Luminex Corporation, Texas, USA) with a laser emitting at 488 nm. The cytometer was calibrated each day of the experiment using the Guava^®^ EasyCheck™ kit, and settings were adjusted before each measurement. The flow rate was set to 1.18 μl.s^−1^, for 30s or until the number of counted events reached 5,000. Data processing was carried out using Guava^®^ InCyte Software version 3.3, from which we exported list-mode data files of measurements on the logarithmic scale. Non-algal particles and dead algae were excluded from the analysis according a cytogram of emissions at Red-B (695/50 nm) and Yellow-B (583/26 nm) band pass filters, enabling a clear discrimination between algae populations and other events thanks to chlorophyll auto-fluorescence (Rescan *et al*. 2020). We also discriminated doublets (i.e. single events that actually consists of two independent cells) from singlets according to the width of the electronic pulse measurement (FSC-W) (Wersto *et al*. 2001). For events categorized as single alive *D. salina* cells, we specifically assessed the environment-specific cell morphology using the Forward Scatter (FSC), Side Scatter (SSC) and fluorescence emission at 695/50 nm band pass filter (Red-B) values as proxy for cells size, complexity (granularity, cytoplasmic contents) (Adan *et al*. 2017), and chlorophyll production (Papageorgiou 2004), respectively. The density of the medium is likely to alter the light signal. To control for this effect, we subtracted from each individual measurement of FSC, SSC and Red-B the mean value of the same parameter values among 1,000 Guava^®^ EasyCheck™ calibrating beads placed in artificial seawater at the same salinity ([NaCl] = 0.8 M, 2.4 M or 4.0 M). We confirmed that traits we measured using FCM closely matched cell size, shape and cellular contents including chlorophyll production, as there was 90.06% (*P* < 0.001) correspondence between PCAs performed with data from FCM vs image processing of epifluorescence microscopy (Procrustes analysis; Supp. Fig. S1). Finally, for each days and salinity, we determined population densities from the ratio of the count of events identified as alive *D. salina* cells to the total volume of acquisition.

### Statistical analyses

#### Population dynamics

For each population and salinity, we calculated the *per-capita* growth rate of the population per day during the phenotypic plasticity assay, as 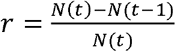, where *N* is the population density (cells × ml^−1^), and *t* a time point of measurement (in days). For *N*(0), we used the initial density of 2 × 10^4^ cells.ml^−1^ that we inoculated in each environment following the acclimation step. We describe days when *r* is highest as the exponential growth phase, while with slower positive growth rates before and after the exponential phase are described as lag and stationary phases, respectively.

#### Morphological variation and dynamics

Flow cytometry allows analyzing thousands of individual cells in seconds, achieving high-throughput phenotyping. Here, the low population density following transfer to a new environment (Days 1-2) limited the number of cells that could be measured by the flow cytometer, accounting for our measurement parameters. To keep a balanced design, we therefore randomly selected up to 150 events categorized as alive *D. salina* per conditions – i.e. population, environment and day –, totalizing more than 2 × 10^5^ individual cells for this study.

We then used a multivariate approach based on Redundancy Analyses (RDA) (Borcard *et al*. 1992) to assess the proportion of morphological variation explained by different predictors. For each *D. salina* cell, we used the morphological measurements (FSC, SSC and Red-B cytometer values) as the multivariate response variable, and ancestral strain identity (CCAP 19/12 or CCAP 19/15), salinity during the plasticity assay, time point of measurement (day), and associated *per-capita* growth rate, as explanatory variables. Variation partitioning was performed separately for populations that evolved in fluctuating *vs* constant environments. For populations that evolved in fluctuating environments, the predictability of environmental change during long-term evolution was included as an additional explanatory variable. As mentioned above, different time series within an autocorrelation treatment may vary in their realized autocorrelation, because of the randomness of the stochastic process in finite time. We therefore computed the realized environmental autocorrelation *ρ* as the correlation between salinities at two subsequent transfers, throughout each salinity time series. We then assessed the environmental predictability as *ρ^2^*, and used it as a continuous explanatory variable.

We quantified the effects of all explanatory factors or variables and their interactions through their contributions to total variation using a multivariate version of *R^2^*, and tested the significance of each *R^2^* by ANOVA-like permutation tests, using 999 randomizations of the data (Borcard *et al*. 1992; Legendre & Legendre 1998; Peres-Neto *et al*. 2006). A significant *salinity* × *predictability* interaction characterized the evolution of plasticity in response to our predictability treatment, a significant *day* × *salinity* interaction characterized a salinity-specific ontogenic trajectory of morphology, and a significant three wise *day* × *salinity* × *predictability* interaction indicated that the evolution of plasticity had different effects at different days along the ontogenic trajectory.

To illustrate the temporal changes in cells morphology following osmotic stress, we represented the mean cell morphology for each day in a morphospace for each targeted autocorrelation 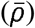. To do so, we first performed a Principal Component Analysis (PCA) of the morphological measurements of the entire dataset, and calculated the centroid for each conditions (i.e. for a given salinity, day and population). We then represented the mean morphology with their standard errors, per day and salinity, by averaging over all populations for each targeted 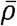, and represented them along the two first PCA axes (Fig. 3B).

#### Evolution of the degree of plasticity

To assess how the magnitude of phenotypic plasticity evolved in our experiment, we first computed the Euclidian distance between the centroids (multivariate means) of each experimental population measured at high (4M) *vs* low (0.8M) salinity. We then compared this degree of plasticity among the experimental populations, to test whether it evolved according to environmental predictability. Specifically, for each day following transfer to the new salinity, we regressed the Euclidian distance of plastic change against *ρ^2^*, as index for environmental predictability. The significance of this regression was assessed by hierarchical non-parametric bootstrapping. We first resampled with replacement 32 populations among the 32 populations that evolved in fluctuating environments. For each population in each salinity, we resampled *n* = 150 cells with replacement. We then recomputed the degrees of plasticity of all populations, and regressed them against *ρ^2^*, iterating the full process 1,000 times. The proportion of simulations with slopes larger than 0 was used to assess the significance of the evolution of lower plasticity in populations evolved in less predictable environments.

We performed all statistical analyses in the environment R version 3.5.3 (R Core Team 2019) with the vegan package version 2.5-4 (Oksanen *et al*. 2019) for the multivariate analyses.

## Results

### Evolution of reduced plasticity

We first investigated the plasticity of cells that had approached phenotypic equilibrium, 10 days after transfer to a new salinity. Comparison of samples from low *vs* high salinities revealed a clear signal of plasticity, with a significant effect of salt concentration on cell morphology (Table 1). Cells from high salinity were smaller and contained less chlorophyll than cells from low salinity (Fig. 2A & Supp. Fig. S2A). However this plasticity was not identical in all experimental lines: those that evolved in different environmental predictabilities differed in their plastic responses to salinity (Fig. 2B; significant *salinity* × *ρ^2^* interaction in Table 1). The magnitude of plasticity, as quantified by the Euclidian distance of phenotypic change between low and high salinities (length of black segments in Fig. 2B, & Supp. Fig. S2B), was positively correlated to the environmental predictability during experimental evolution, with lines from less predictable environments displaying reduced plasticity (linear regression: *R^2^* = 0.621, *P* < 0.001; Fig. 2C). This result was replicated over the two ancestral strains (Fig. 2C) despite their differences on other features such as their population dynamics (Fig. 3A & Supp. Fig. S3), and also held for isogenic populations (Fig. S4A). The latter confirmed that the observed morphological differences between salinities were the result of phenotypic plasticity, rather than rapid selection of salinity-specific genotypes over the assay experiment (since isogenic populations display no genetic variation).

**Table 1.**
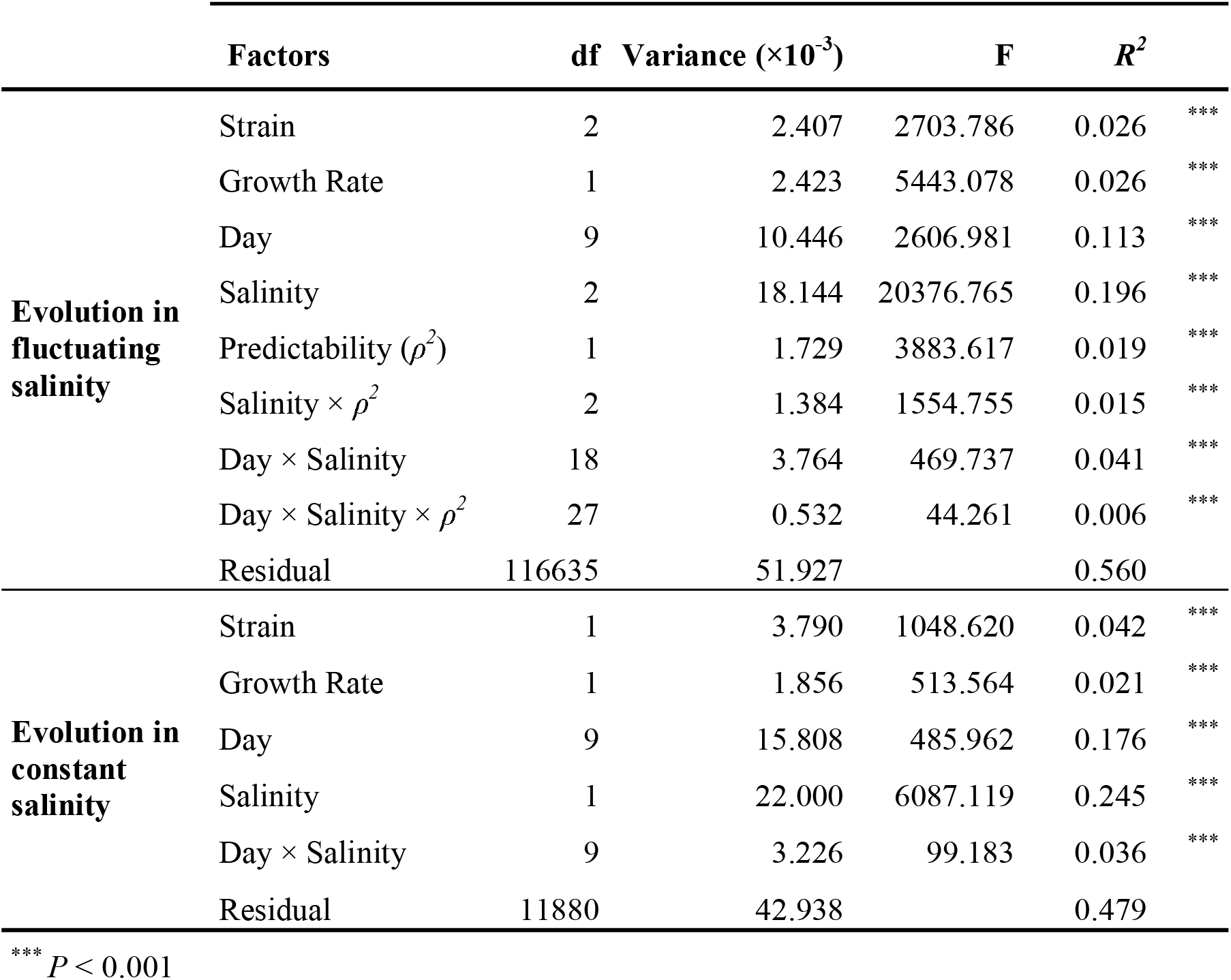
Partitioning of cell morphological variation. The effect of all explanatory factors and their interactions on multivariate patterns of cellular variation are quantified by their *R^2^* (proportion of total variation explained), and the significance of each *R^2^* was tested by ANOVA-like permutation tests using 999 randomizations of the data (Legendre & Legendre 1998).

| | Factors | df | Variance ($\times 10^{-3}$) | F | $R^2$ | |
| --- | --- | --- | --- | --- | --- | --- |
| **Evolution in fluctuating salinity** | Strain | 2 | 2.407 | 2703.786 | 0.026 | *** |
|  | Growth Rate | 1 | 2.423 | 5443.078 | 0.026 | *** |
|  | Day | 9 | 10.446 | 2606.981 | 0.113 | *** |
|  | Salinity | 2 | 18.144 | 20376.765 | 0.196 | *** |
| | Predictability ($\rho^2$) | 1 | 1.729 | 3883.617 | 0.019 | *** |
| | Salinity $\times \rho^2$ | 2 | 1.384 | 1554.755 | 0.015 | *** |
| | Day × Salinity | 18 | 3.764 | 469.737 | 0.041 | *** |
| | Day × Salinity $\times \rho^2$ | 27 | 0.532 | 44.261 | 0.006 | *** |
|  | Residual | 116635 | 51.927 |  | 0.560 |  |
| **Evolution in constant salinity** | Strain | 1 | 3.790 | 1048.620 | 0.042 | *** |
|  | Growth Rate | 1 | 1.856 | 513.564 | 0.021 | *** |
|  | Day | 9 | 15.808 | 485.962 | 0.176 | *** |
|  | Salinity | 1 | 22.000 | 6087.119 | 0.245 | *** |
| | Day × Salinity | 9 | 3.226 | 99.183 | 0.036 | *** |
|  | Residual | 11880 | 42.938 |  | 0.479 |  |

\*\*\* $P < 0.001$

**Fig. 2.**
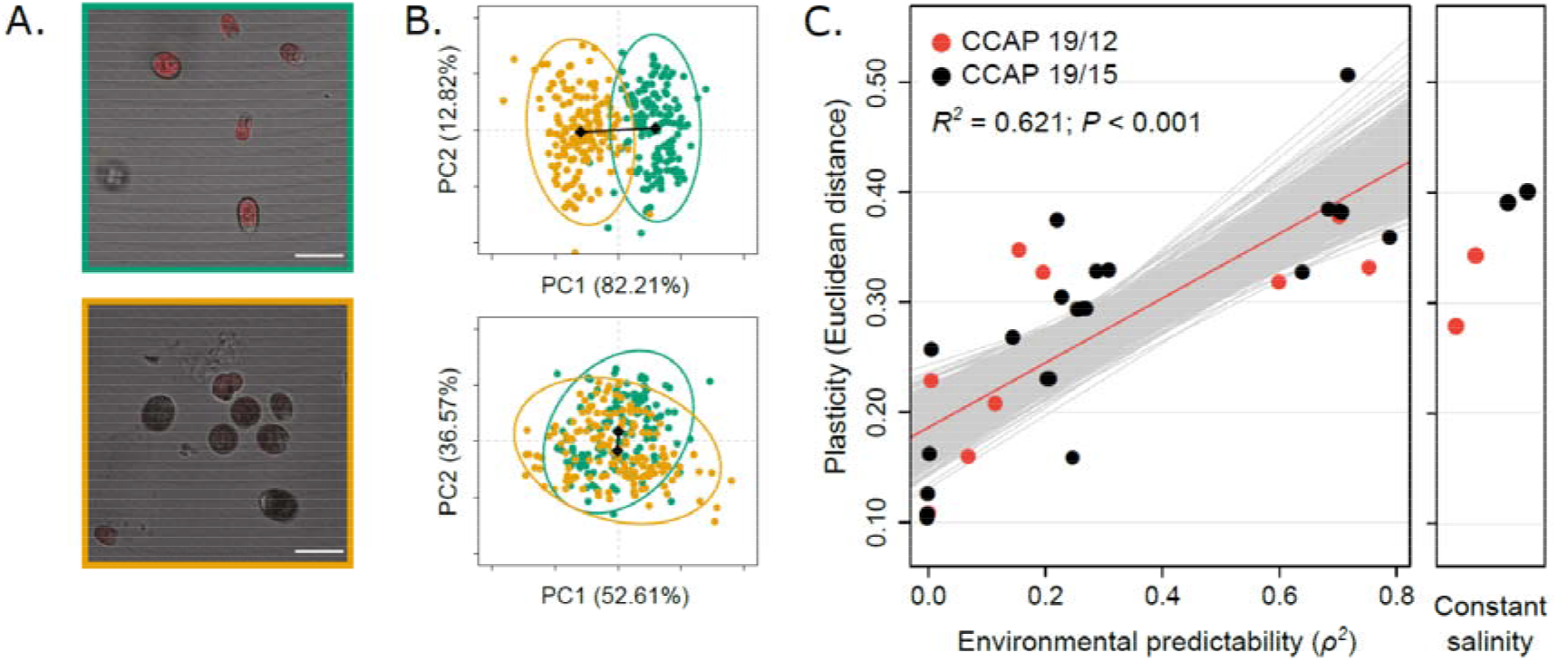
Evolution of morphological plasticity in response to environmental predictability. A. Example of salinity-specific cell morphologies. Red zones in composite images represent chlorophyll fluorescence intensity and white bars give the scale (20μm) for populations in low (green) and high (orange) salinities. B. Morphological variation of cells after 10 days in low (green) and high (orange) salinities for populations that have evolved under predictable (upper graph) *vs* unpredictable (lower graph) environmental fluctuations. C. Evolution of the degree of plasticity at day 10. Standard error based on 1000 bootstraps are also plotted for each dot, but not visible. Red line is the regression slope, and the grey lines show 1000 regression slopes calculated on bootstrapped samples.

**Fig. 3.**
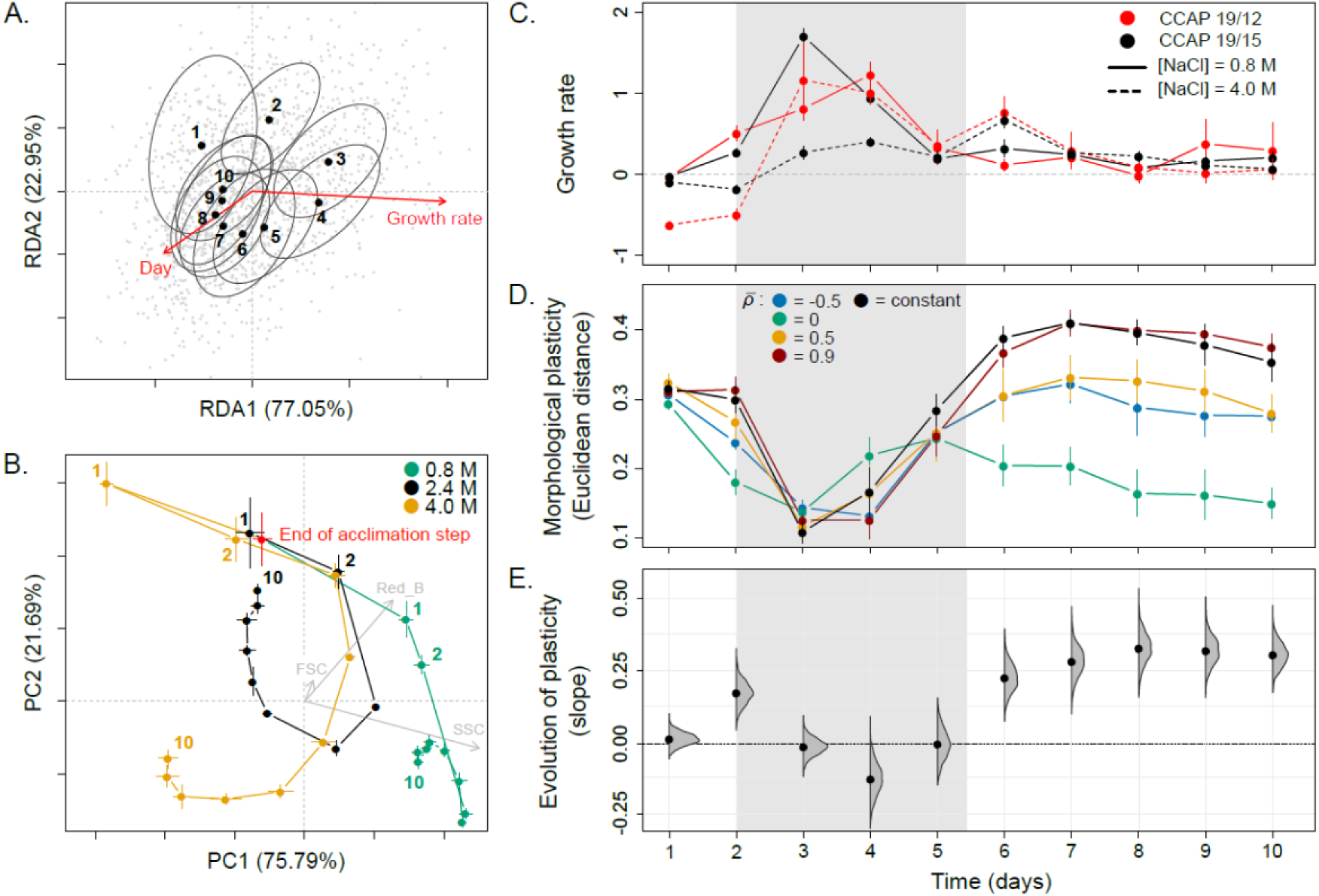
Evolution of plasticity along the ontogenic trajectory. A. The ontogenic trajectory of cell morphology for an isogenic population transferred to fresh medium with unchanged salinity is shown as centroids (dots) and covariance matrices (ellipses) for each day. Arrows indicate the RDA loadings. B. Trajectories of four isogenic populations from the same evolved line in three different salinities. PC loadings of flow cytometry measurements (grey arrows) were rescaled by dividing them by 20 to facilitate graphical representation. C. Per capita population growth rate per day, across all evolved lines. D. Temporal variation in the degree of phenotypic plasticity of the four treatments of expected long-term temporal autocorrelation 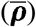, and populations that evolved in constant salinity (black). E: Evolution of plasticity along the ontogeny. The mean slope of the regression of the degree of plasticity (Euclidian distance between environments) against environmental predictability (*ρ^2^*) is shown for each day post transfer to a new salinity. Density plots represent slopes estimated on 1000 bootstrapped samples of the raw data (same as grey lines in Fig. 2C for day 10). For C to E, the grey dashing delimits the phase with highest growth rate. Means with their standard errors are represented as dots and error bars respectively.

We also maintained four lines at constant salinity ([NaCl] = 2.4M) throughout the experiment, as controls for the influence of environmental fluctuations. Lines that evolved in constant environments (right panel in Fig. 2C & Supp. Fig. S4A) had a similar degree of plasticity as lines evolved under highly predictable environmental fluctuations (*P* = 0.295). This suggests that unpredictable environments exerted a stronger selective pressure against plasticity (Gavrilets & Scheiner 1993; Lande 2009) than any putative cost associated to the maintenance of plasticity in a constant environment (DeWitt *et al*. 1998). Interestingly, we found similar degrees of plasticity between populations that evolved in treatments with autocorrelations 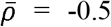 and 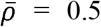 (*P* = 0.867), which have the same expected predictability of changes (*ρ^2^* = 0.25), but different magnitude of transitions upon each transfer (Supp. Fig. S5).

### Ontogeny of evolved plasticity

We then turned to the entire morphological trajectory over 10 days, to investigate how the evolution of plasticity unfolds along developmental time scales. When cells were transferred at low density (~2 × 10^4^ cells.ml^−1^) to fresh medium with unchanged salinity, their morphology followed a loop-shaped trajectory over 10 days, with initial and final phenotypes markedly differing from phenotypes at intermediate times (Fig. 3A). These changes concerned cell size and granularity for the first days (up to days 2-3: bigger cells when reaching the exponential phase), followed by changes in chlorophyll content (days 3-4 to 10: decreasing chlorophyll content in the stationary phase) (Fig. 3A-B & Supp. Fig. S1A). As this temporal trend in morphology was also found for isogenic populations with effectively no opportunity for natural selection (Fig. 3A-B), it does not reflect genetic changes in the population. Instead, these changes can be described as ontogenic, in line with the extended definition of ontogeny/development as a sequence of cellular states, applying across unicellular and multicellular organisms (Gilbert 2000). Part of this ontogenic morphological variation was explained by population growth rate (Table 1), an aggregate population-level outcome of life-history traits of individual cells (division and death rate), which is typically used as an indicator of physiological state in microbiology (Maharjan *et al*. 2013). Focusing on populations maintained in the same salinity as during acclimation ([NaCl] = 2.4 M), we detected a clear effect of growth rate (*R^2^* = 0.046, *P* < 0.001) on morphological variation (red arrow in Fig. 3A; same effect for both ancestral strains, *growth rate × strain* interaction: *P* = 0.504), independent from salinity changes. However, the population growth rate was not sufficient to explain the ontogenic trajectory, and there remained a significant marginal effect of the time spent in the new environment (*R^2^* = 0.031, *P* < 0.001).

Ontogenic trajectories also differed between salinities. Upon transfer to a new salinity, morphologies in the hypo-*vs* hyper-osmotic environments first rapidly diverged in opposite directions from the acclimated morphology, 4h after salinity change (day 1 in green and yellow vs red dot; Fig. 3B), before converging during the exponential phase, and finally diverging again to salinity-specific morphologies over the stationary phase (Fig. 3B & Supp. Fig. S2A). This is consistent with known responses of *D. salina* to salinity change, which first involve an immediate – sometimes drastic – change in cell volume caused by water intake/loss, followed by slower accumulation of salt-induced proteins and metabolites (notably via changes in gene expression (Chen & Jiang 2009; Fang *et al*. 2017)), which restore cell shape and eventually leads to long-term changes in cell content such as lipid, carotene, and glycerol accumulation (Ben-Amotz *et al*. 2009). As a result of these differences in morphological trajectories between salinities, the plasticity of cell morphology was temporally variable (significant *day × salinity* interaction; Table 1). Plastic differences in morphology also followed a loop over 10 days, where initial and final differences diverged from those at intermediate times (Supp. Fig. S2B). The degree of plasticity was highest at low population growth rates characteristic of the lag and stationary phases, and lowest during the exponential phase (significant *salinity × growth rate* interaction, Table 1, Fig. 3C-E and Supp. Fig. S2).

Evolution of plasticity in response to environmental predictability had different impacts at distinct stages of the ontogenic trajectory. We found a positive relationship between the level of plasticity and environmental predictability during phases of slow population growth (at day 2, and from days 6 to 10; Fig. 3C-E), but no relationship at day 1 and during the exponential phase (days 3 to 4; Fig. 3C-E; similar results were observed for isogenic populations, Supp. Fig. S4B). That there was little evolution of plasticity during the exponential phase is consistent with the finding that morphological plasticity is generally lowest in this phase (Fig. 3C-D). In contrast, plasticity is high shortly after salinity transfer (day 1, Fig. 3D), but mostly because of passive, reflex responses to osmotic stress, which are certainly less prone to evolution than longer term physiological responses involving specific gene expression and production of metabolites and chlorophyll.

## Discussion

We have shown that phenotypic plasticity, a major component of phenotypic change in the wild (Scheiner 1993; Schlichting & Pigliucci 1998; West-Eberhard 2003; Pelletier *et al*. 2007; Ozgul *et al*. 2010; Ellner *et al*. 2011), and a key mechanism for population persistence in a changing environment (Chevin *et al*. 2010; Reed *et al*. 2010; Chevin *et al*. 2013b; Vedder *et al*. 2013; Ashander *et al*. 2016; Phillimore *et al*. 2016), can evolve experimentally in response to environmental predictability, in the direction predicted by theory (Gavrilets & Scheiner 1993; de Jong 1999; Lande 2009; Botero *et al*. 2015; Tufto 2015). This plasticity concerned morphological traits that were previously described to be involved in salinity tolerance in *Dunaliella salina* (Azachi *et al*. 2002; Oren 2005; Ben-Amotz *et al*. 2009), a species for which we have shown that plastic responses to past environments can largely drive population dynamics and extinction risk in a randomly fluctuating environment (Rescan *et al*. 2020).

Contrary to common practice in experimental evolution with microbes (Elena & Lenski 2003; Buckling *et al*. 2009), we have assayed evolved changes by measuring multiple individual traits (rather than aggregate population traits) across environments and over time. This approach, tending towards phenomics, allowed us to describe an ontogenic sequence of cell morphology, consistent with osmotic response mechanisms operating at different time scales. These responses ranged from immediate physical change in cell shape (passive plasticity) occurring within the first few seconds/minutes, to long-term physiological regulations involving changes in gene expression (active plasticity), which usually start within 12-24h (Chen & Jiang 2009; Fang *et al*. 2017). These consecutive morphological states of cells, which we here described as an ontogenic sequence (Gilbert 2000), can also be interpreted as alternative phenotypes favored at different population densities (*r* vs *K*-selection (MacArthur 1962; Sæther *et al*. 2016)), or plastic responses to environmental variables other than salinity that change along time in a batch culture (e.g. resource abundance and quality) (Collot *et al*. 2018).

Evolution of active plasticity in response to environmental predictability was essentially restricted to physiological states associated with phases of slow – or even null – population growth in our experiment. Population growth status, indicative of the physiological state of individual cells, is known to be associated with multiple phenotypic traits micro-organisms (Maharjan *et al*. 2013). The lack of plasticity during the exponential phase likely results from a particular morphology associated to rapid cell division, which masks the influence of salinity. However, the late expression (day 5-10) of morphological changes that we observed in response to osmotic stress was initiated as soon as the start of the active plasticity (day 2). Similarly, we also observed the evolution of reduced plasticity according to environmental predictability as early as the beginning of the ontogenic sequence of cells morphology.

Our results demonstrate that long-term experimental evolution under complex, ecologically realistic patterns of environmental variation, coupled with intense high-throughput phenotyping at the individual level, allows testing fundamental predictions in evolutionary ecology that are barely approachable in natural settings. Our finding that phenotypic plasticity can evolve in response to environmental predictability proves important for the prospects for population persistence in the face of global warming, and other anthropogenic changes. Indeed these changes consist not only of trends in mean environments, but also alterations in the magnitude and predictability of natural environmental fluctuations (Wigley *et al*. 1998; Boer 2009). Theoretical work has made it clear that phenotypic plasticity can strongly influence extinction risk in response to changing environmental predictability (Reed *et al*. 2010; Chevin *et al*. 2013a; Botero *et al*. 2015; Ashander *et al*. 2016), which was recently confirmed empirically using laboratory experiments (Proulx *et al*. 2019; Rescan *et al*. 2020). In particular, this theory has shown that evolution of lower phenotypic plasticity can reduce extinction risk under reduced environmental predictability, by decreasing the magnitude of mismatches between the population mean phenotype and the optimum phenotype determined by the environment (Chevin *et al*. 2013a; Ashander *et al*. 2016). Our demonstration that such evolution of reduced plasticity can indeed occur over a few hundred generations indicate that this may be an important mechanism by which species may persist under climate change.

## Acknowledgements

We are grateful to S. Bedhomme, P. Nosil and R. Villoutreix for comments on an earlier version of the manuscript. We thank the MRI-IGMM and MRI-CRBM platforms (Montpellier) for access to the cell sorter and microscopy, respectively. This work was supported by the European Research Council (Grant 678140-FluctEvol) to L-MC, and a Fonds de Recherche du Québec - Nature et Technologies (FRQNT) fellowship to CL.

## Supplementary Information

Supplementary information include Supplementary Material 1, and Supplementary figures S1 to S5.

## Supplementary Material 1

Comparison of flow cytometry vs confocal microscopy methods to assess cells morphology

We carried out microscopy images analyses on a small subset of populations, to confirm the accuracy of flow cytometry for assessing cell morphology.

## Methods

As for the main plasticity assay, we first acclimatized four randomly chosen evolved populations to the mean salinity ([NaCl] = 2.4 M) during 10 days. We then transferred each of them to low ([NaCl] = 0.8 M) and high ([NaCl] = 4.0 M) salinity for 10 others days. We finally measured cell morphology using flow cytometry and dragonfly confocal microscopy.

### Dragonfly confocal microscopy

We sampled ~40μL of cultures from each condition (*n* = 8, for 4 populations in 2 salinities), and scanned them on a Dragonfly (COM4) spinning disk confocal microscope with 2 cameras EMCCD iXon888 Life (Oxford Instrument - Andor). A total of 100 images was taken under both transmission and confocal fluorescence modes, to assess cells shape and chlorophyll content, respectively. Wavelengths excitation and emission were at 488 nm and 700/775 nm, respectively, for confocal fluorescence.

We then used the automatic image analysis software Fiji (Schindelin *et al*. 2012) to characterize cells morphology. First, we used the function *Analyze Particles* to isolate cells in each picture, and manually validate each detected particle as an actual *D. salina* cell. For each particle, we then recorded morphological descriptors including the area and aspect ratio (AR: major/minor axis), the average gray - sum of the gray values of all the pixels in the selection divided by the number of pixels (Grey_Int and Grey_sd for mean and standard error respectively) - for the transmission mode, and fluorescence intensity (Fluo_Int and Fluo_sd for mean and standard error respectively) within the selection.

### Flow cytometry

We passed a 150 μL sample of each condition through flow cytometry, using the same method as described in the main text. We then randomly selected 30 events characterized as alive *Dunaliella salina* cells per condition, to keep a balanced sampled size with the microscopy dataset. We then used FSC, SSC and Red-B values, followed by light signal correction using Guava^®^ EasyCheck™ kit calibrating beads, to assess cells morphology.

### Statistical analyses

We tested the concordance of the morphospaces obtained by using the two methods, as proposed by Wang *et al*. (2010). We first carried out independent principal component analyses of morphology for flow cytometry and microscopy. We thereafter extracted the centroids on PCA plot for each group of cells, characterized by their population and salinity. Finally, we aligned the centroids’ PCA coordinates of flow cytometry morphospace with the microscopy ones using a Procrustean superimposition approach (Gower 1975; Legendre & Legendre 1998). The degree of concordance between the ordinations was assessed with a correlation-like statistic *r-*PROTEST(Jackson 1995), derived from the symmetric sum of squares, where 1 and 0 denote a perfect vs totally absent concordance, respectively. The significance of this correlation was tested by permutation, using 999 randomizations against the null hypothesis that there is no concordance between the two ordinations (Jackson 1995).

---

**Supp. Fig. S1.**
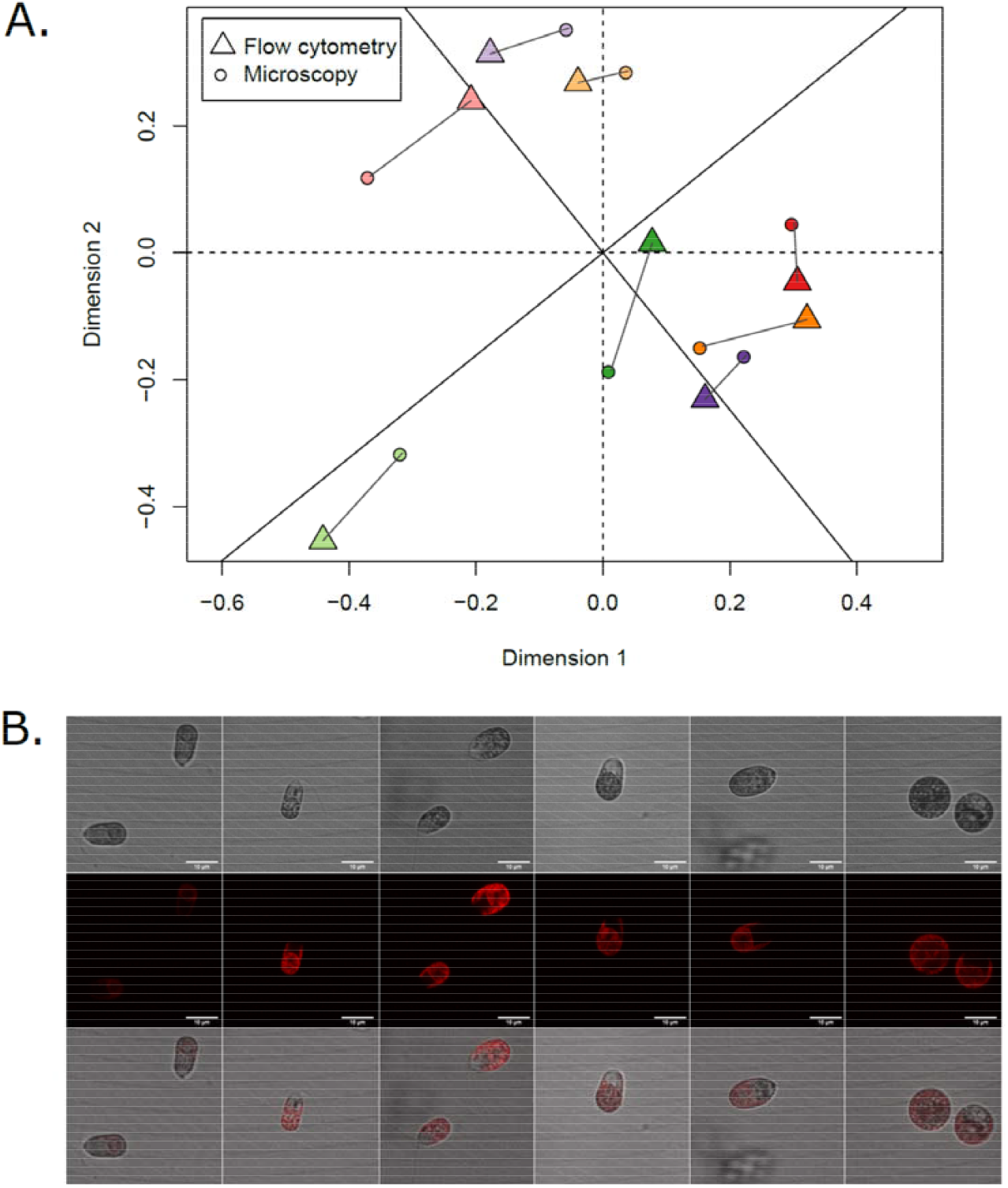
Comparison of flow cytometry vs confocal microscopy methods to assess cells morphology. A. Procrustes errors between PCAs performed on flow cytometry (triangle) and confocal microscopy (circle) parameters. Each color represent the centroids of a given population, with lighter vs darker colors for low (0.8 M) vs high (4.0 M) salinity, respectively. B. Example of cell morphology. Images were taken using a dragonfly confocal microscope, under the transmission (first line) and confocal in fluorescence (second line) acquisitions respectively, and a composite of both acquisitions (third line, similar to Fig. 2B in the main text). White scale bar represents 10μm.

**Supp. Fig. S2.**
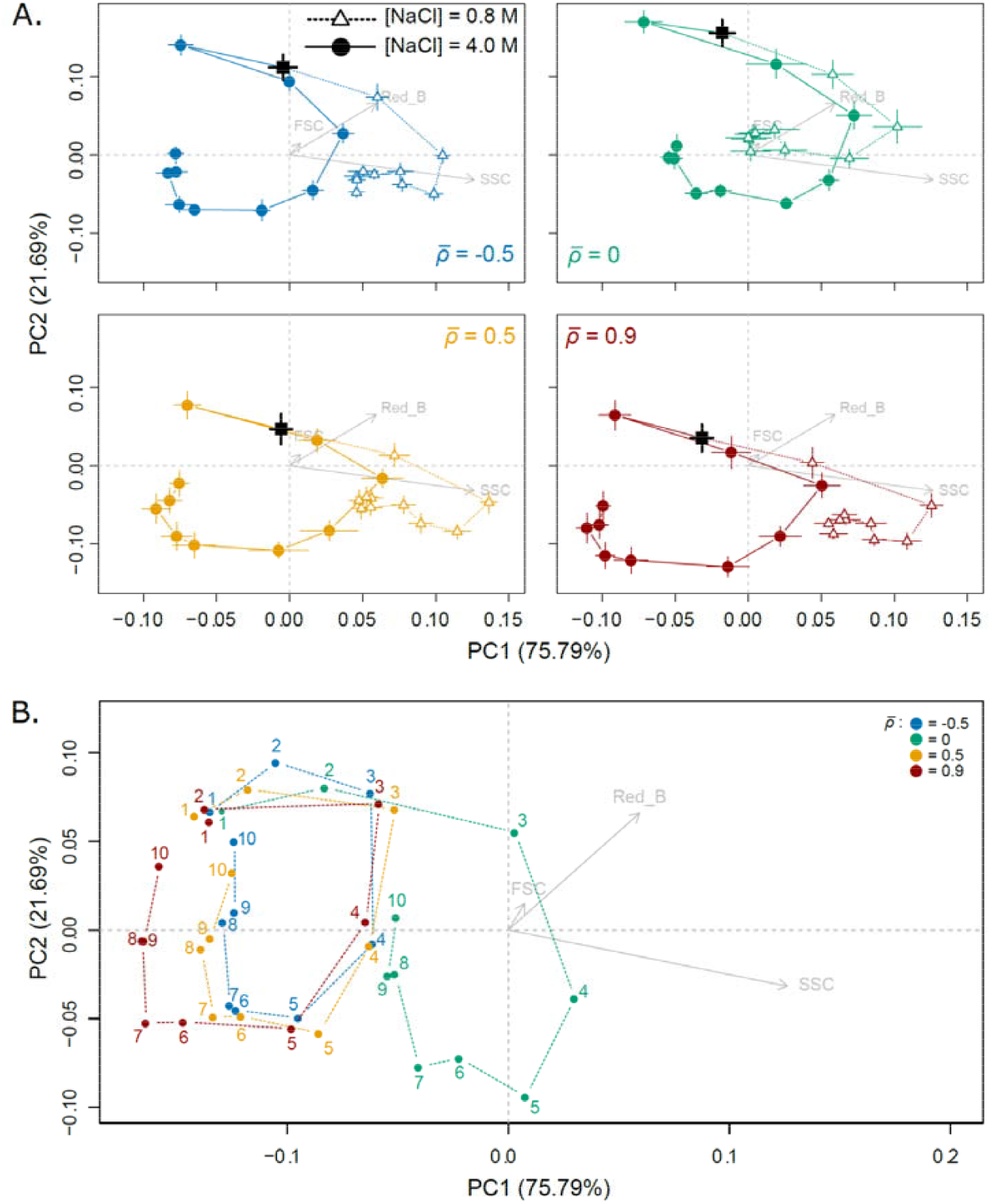
Ontogenetic trajectories and dynamics of plasticity over time. A. Morphological trajectories in two salinities for each autocorrelation treatment 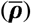. Mean cell morphologies with their standard errors, per day and salinity, are calculated by averaging over lines belonging to the same targeted 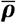, and represented along the two first PCA axes. Vectors of PC loadings of response variables (FSC, SSC and Red_B grey arrows) were divided by 20 to facilitate graphical representation. Black squares represent cell morphology at the end of acclimation step. B. Trajectories of plasticity over time. Morphological differences between salinities for each treatments were computed as vector at high salinity *minus* the one at low salinity, for each day (number).

**Supp. Fig. S3.**
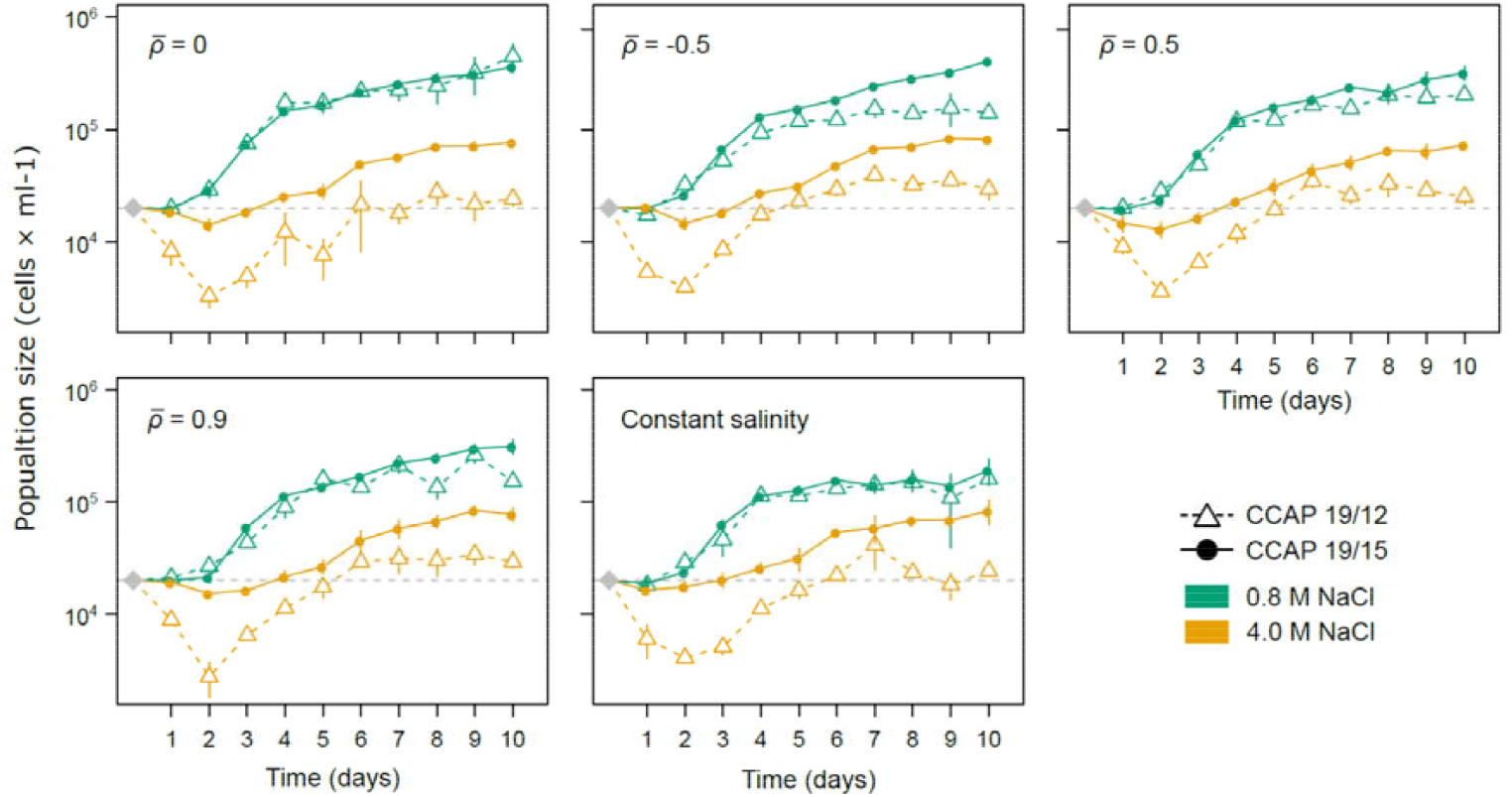
Population dynamics in low and high salinities. For each evolutionary treatment (temporal autocorrelation 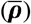 or constant salinity at [NaCl] = 2.4M), mean population size (cells × ml^−1^) with their standard errors, per day and salinity, were calculated by averaging over lines belonging to the same strain. Grey diamond is the expected initial population size (2 × 10^4^ cells × ml^−1^) inoculated in the new salinity at the beginning of the plasticity assay, and day 1 was measure 4h after this salinity change.

**Supp. Fig. S4.**
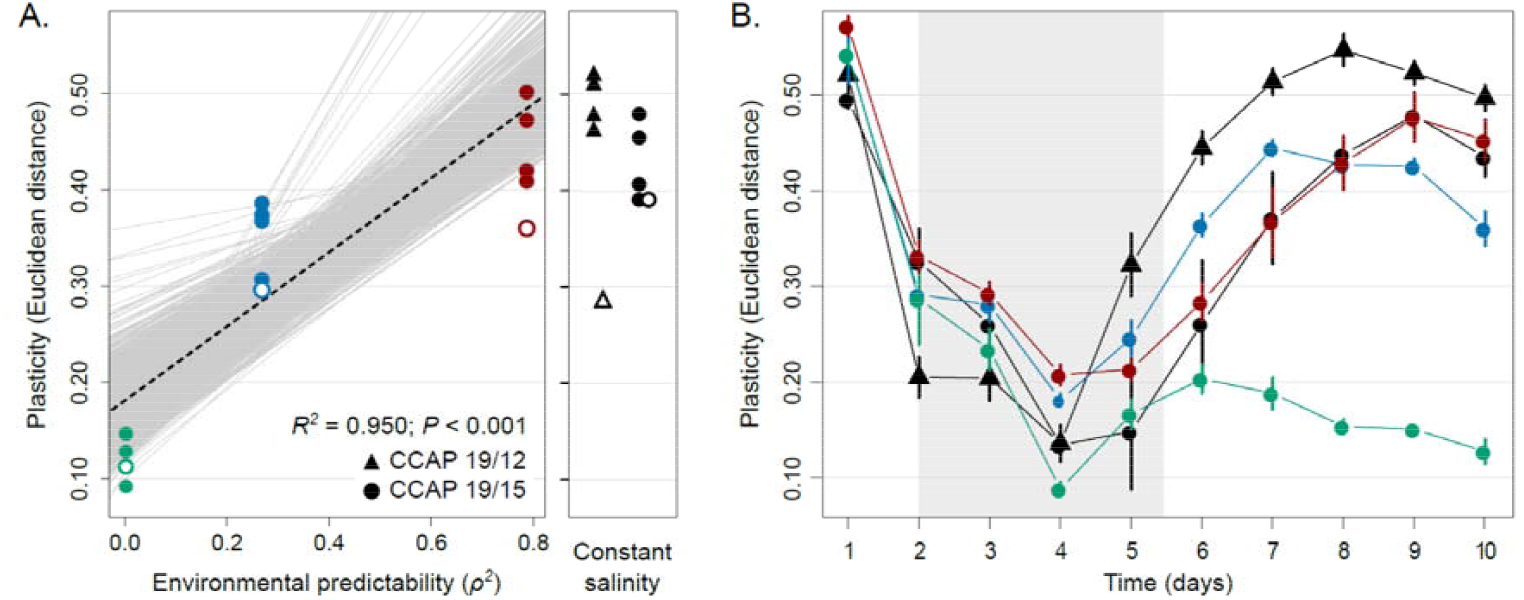
Analyses of isogenic populations. A. Evolution of plasticity at day 10. Isogenic populations were founded from a single cell from tree different evolved lines under *ρ* = −0.022 (green), *ρ* = −0.522 (blue) and *ρ* = 0.889 (red) and two strain (CCAP 19/12 as triangle and CCAP 19/15 as circle) from constant environment. The degree of plasticity of populations, measured as the Euclidian distance of mean cell morphology between low and high salinities, is plotted against the predictability *ρ^2^* of environmental fluctuations of these populations that have experienced during experimental evolution. Standard error based on 1000 bootstraps are also plotted, but not visible. The regression slope is represented as red line, and the grey lines encompass 1000 regression slopes calculated for each bootstrap. The panel on the right shows plasticity in populations that evolved under constant salinity, corresponding to the mean of fluctuating treatments. Variation partitioning revealed significant morphological differences among isogenic populations founded from a given evolved lines (*R^2^* = 0.009; *P* < 0.001), but those remained smaller than morphological variation among different evolved lines (*R^2^* = 0.051; *P* < 0.001), when controlling for the effect of salinity, day, and their interaction on the total variation. Filled and open symbols represent isogenic populations and the evolved lines they come from, respectively. B. Temporal variation in the degree of phenotypic plasticity. For each population, Euclidean distances were computed between mean cell morphologies in [NaCl] = 0.8M vs 4.0M. Means and standard errors were assessed from 4 isogenic populations for each evolved lines.

**Supp. Fig. S5.**
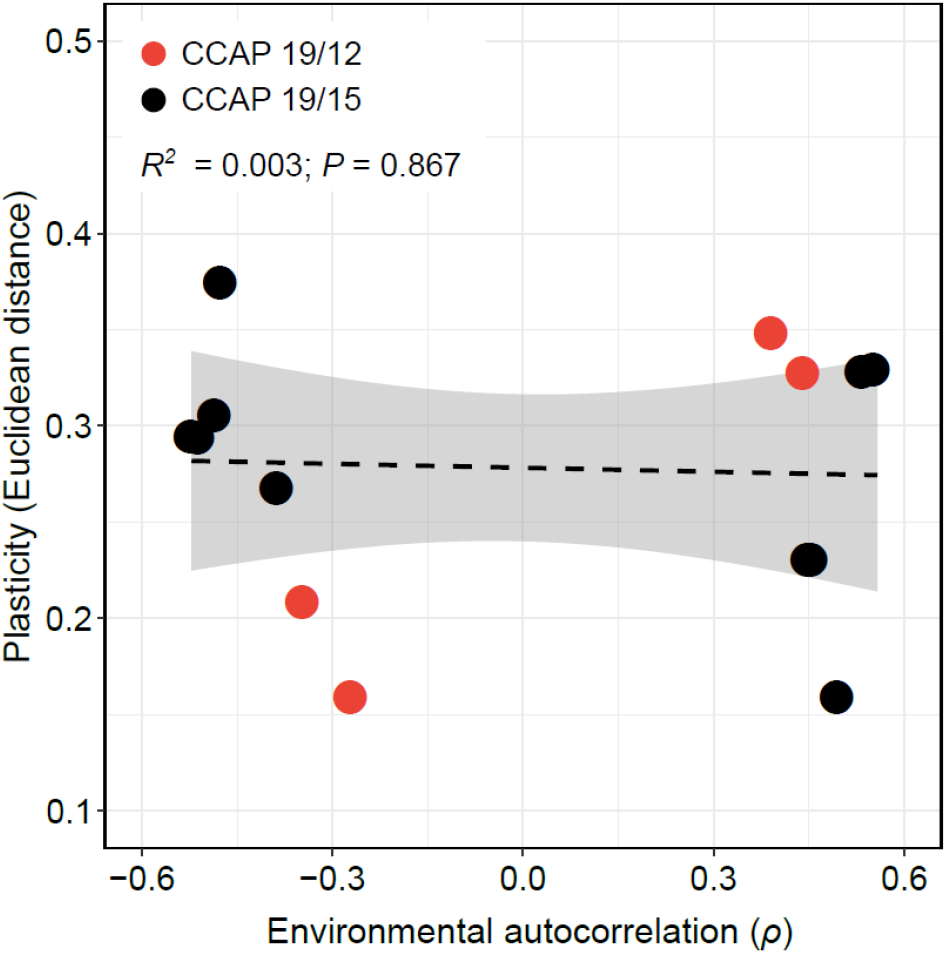
Influence of the magnitude of environmental transitions for a given environmental predictability. The degree of plasticity of populations, measured as the mean Euclidean distance of cell morphology after 10 days in low vs high salinities, is plotted against the environmental autocorrelation 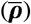. Negative and positive autocorrelation displayed the same environmental predictability 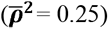, but differed in the intensity of environmental changes between two successive transfers. The regression slope is also represented as black dashed line, and the shaded region shows the 95% CI of this regression.

